# Gut markers of bodily self-consciousness

**DOI:** 10.1101/2021.03.05.434072

**Authors:** Alessandro Monti, Giuseppina Porciello, Maria Serena Panasiti, Salvatore M. Aglioti

## Abstract

Bodily self-consciousness, the state of mind that allows humans to be aware of their own body, forms the backdrop for almost every human experience, yet its underpinnings remain elusive. Here we combine an ingestible, minimally invasive capsule with surface electrogastrography to probe if gut physiology correlates with bodily self-consciousness during a virtual bodily illusion. We discover that specific patterns of stomach and bowel activity (temperature, pressure, pH, and gastric peak frequency) covary with specific facets of bodily self-consciousness (feelings of body agency, location, and disembodiment). These results uncover the hitherto untapped potential of minimally invasive probes to study the link between mental and gut states, and reveal a deep visceral pathway to the self-conscious perception of ourselves as embodied beings.

## Introduction

Humans are ordinarily aware that their body is part and parcel of their self^1–3^. Physiological, clinical, and behavioural data suggest that bodily self-consciousness arises when the central nervous system integrates pieces of body-related information coming from the external and internal senses in a coherent fashion^4,5^. While the contribution of sight, touch, heartbeats and breathing to corporeal awareness is now ascertained^6–9^, the role of deep, sub-diaphragmatic signals is unknown. Despite their potential significance for higher-order cognitive functions^10^, organs located in the abdominal cavity, such as the stomach and the intestine, are difficult to reach and probe without relying on invasive methods^11,12^.

Gut signals are conveyed to the central nervous system via vagal and spinal afferents^13–15^ primarily targeting the somatosensory and insular cortices. In turn, this activity is modulated by top-down control exerted by central and enteric neurons^16,17^. While the gut-brain loop has the clear homeostatic purpose of regulating food intake, it has been proposed that gut signals, once in the brain, may also influence a variety of higher-order processes, including corporeal awareness^18,19^. Gut physiology *per se* is rarely attended to or perceived at the conscious level. Nevertheless, we reasoned that ongoing gut activity could provide the brain with the kind of internal information needed to give us the normal, everyday sensation that our body is not empty.

To shed light on the visceral roots of bodily self-consciousness, we asked healthy male participants (*N* = 31) to ingest a wireless capsule (SmartPill™) fitted with sensors tracking temperature, pressure, and pH values across the entire gastro-intestinal tract in real time^20^. After ingesting the capsule, participants underwent a simplified version of the ‘embreathment’ bodily illusion that we recently discovered^9^. This illusion induces strong feelings of owning, controlling, and dwelling into a virtual body that looks, lays, and breathes as the physical body of the participant (congruent condition) and no such feelings towards a virtual body that does not match the appearance, position, and breathing of the physical body (incongruent condition). During each condition, we also monitored the gastric contractions of our subjects through surface electrogastrography (EGG)^21^ (**Fig**.**1**).

**Figure 1.**
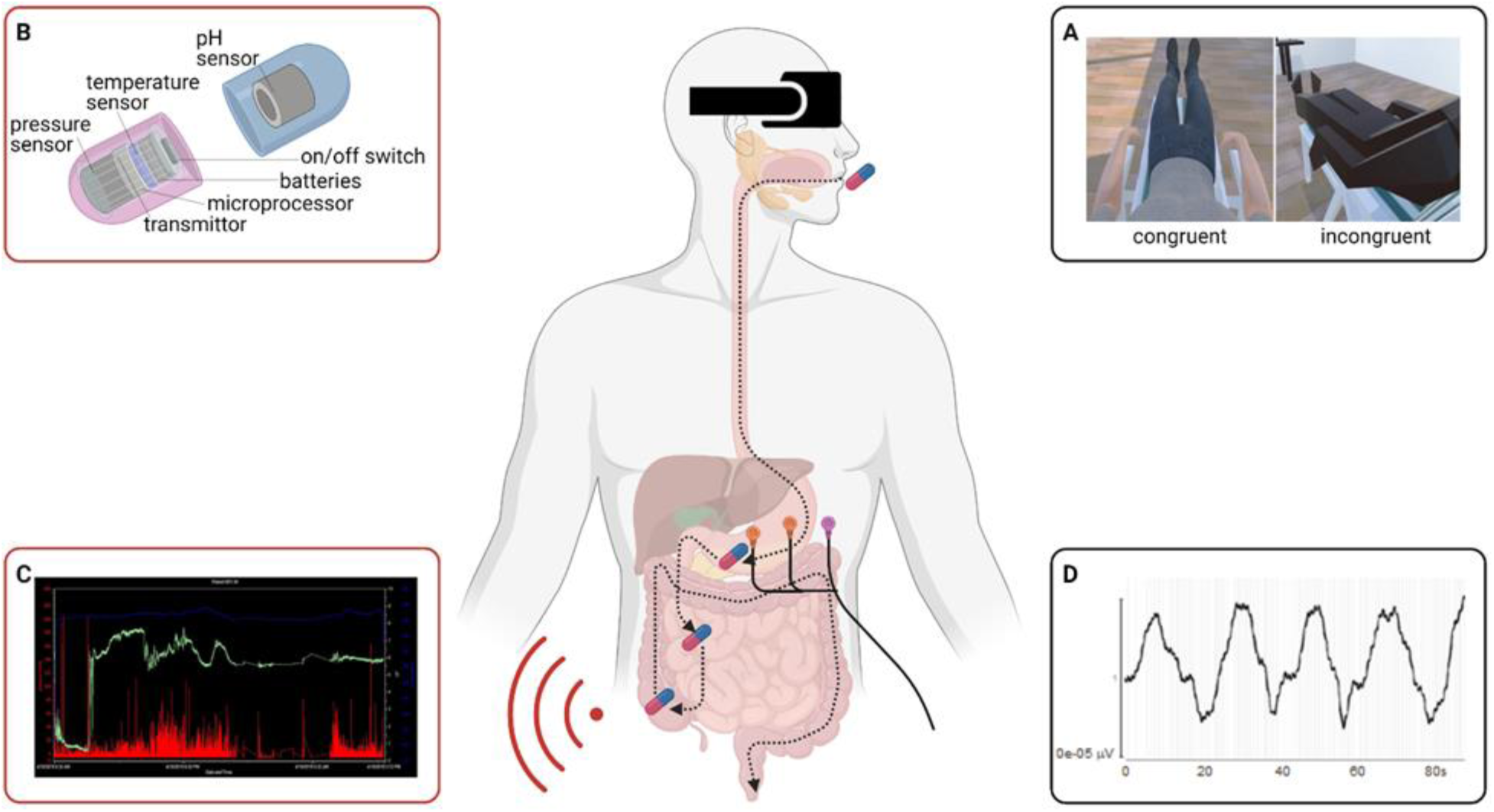
Experimental methods. Participants are administered a simplified ‘embreathment’ bodily illusion (**A**) in which they embody an avatar that looks, lays, and breathes like them (congruent condition) or not (incongruent condition). While they undergo the illusion, their gut is monitored through the SmartPill (capsule icon), which is fitted with a variety of sensors (**B**). This is done three times, as the pill moves down the three main gut segments (stomach, small bowel, and large bowel). The first time, since the capsule lies in the stomach, Smart Pill measures are complemented by electrogastrography (electrode icons). A standard SmartPill graph (**C**) plots temperature (blue line), pH (green line), and pressure (red bars) across the different segments of the gastro-intestinal tract over time. A typical electrogastrographic recording (**D**) shows gastric myoelectric activity generated by interstitial cells of Cajal. Inset (B) is adapted from ^11^.

To assess if and how each district of the gastro-intestinal tract relates to bodily self-consciousness, we administered the illusion thrice: first when the SmartPill was in the stomach, then after the capsule entered the small bowel, and finally when it reached the large bowel (see **Methods** below for details). We hypothesized that objective gut signals recorded during the experimental conditions would predict the subjective feelings of owning, controlling, or dwelling into a body, as gauged through a visual analogue scale (VAS) questionnaire^9^ (see **Table 1**).

**Table 1.**
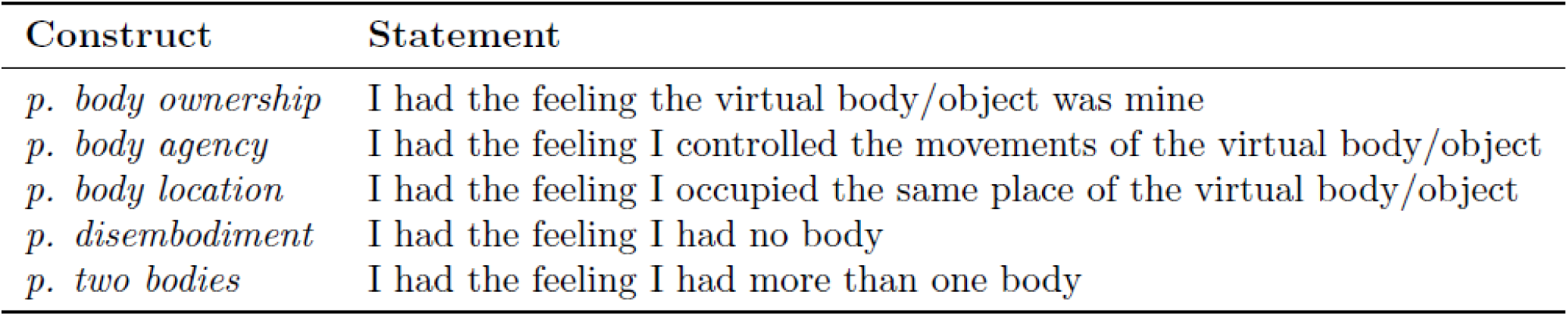
Bodily self-consciousness questionnaire. *p*.: perceived.

## Results

Two linear mixed-effects models measured how specific feelings of bodily self-consciousness induced by our bodily illusion depended on the pH, pressure, and temperature values recorded by the SmartPill (Model 1) and on gastric peak frequencies recorded by surface EGG (Model 2) (see **Methods** for a detailed description of the models and the **Supplementary Information** for a comprehensive report of the results).

### Model 1 (SmartPill) results

#### Sense of disembodiment covaries with gut temperature and pH

We found that the higher the temperature across the gastro-intestinal tract, the less our participants reported to feel disembodied when they observed a virtual body. For each 1°C increase in stomach, small bowel, or large bowel temperature, there was a 13.49 ± 5.94 points decrease in predicted VAS ratings of disembodiment (*t* = −2.272, *p* = 0.0237; **Fig. 2** left and **Table S2**).

**Figure 2.**
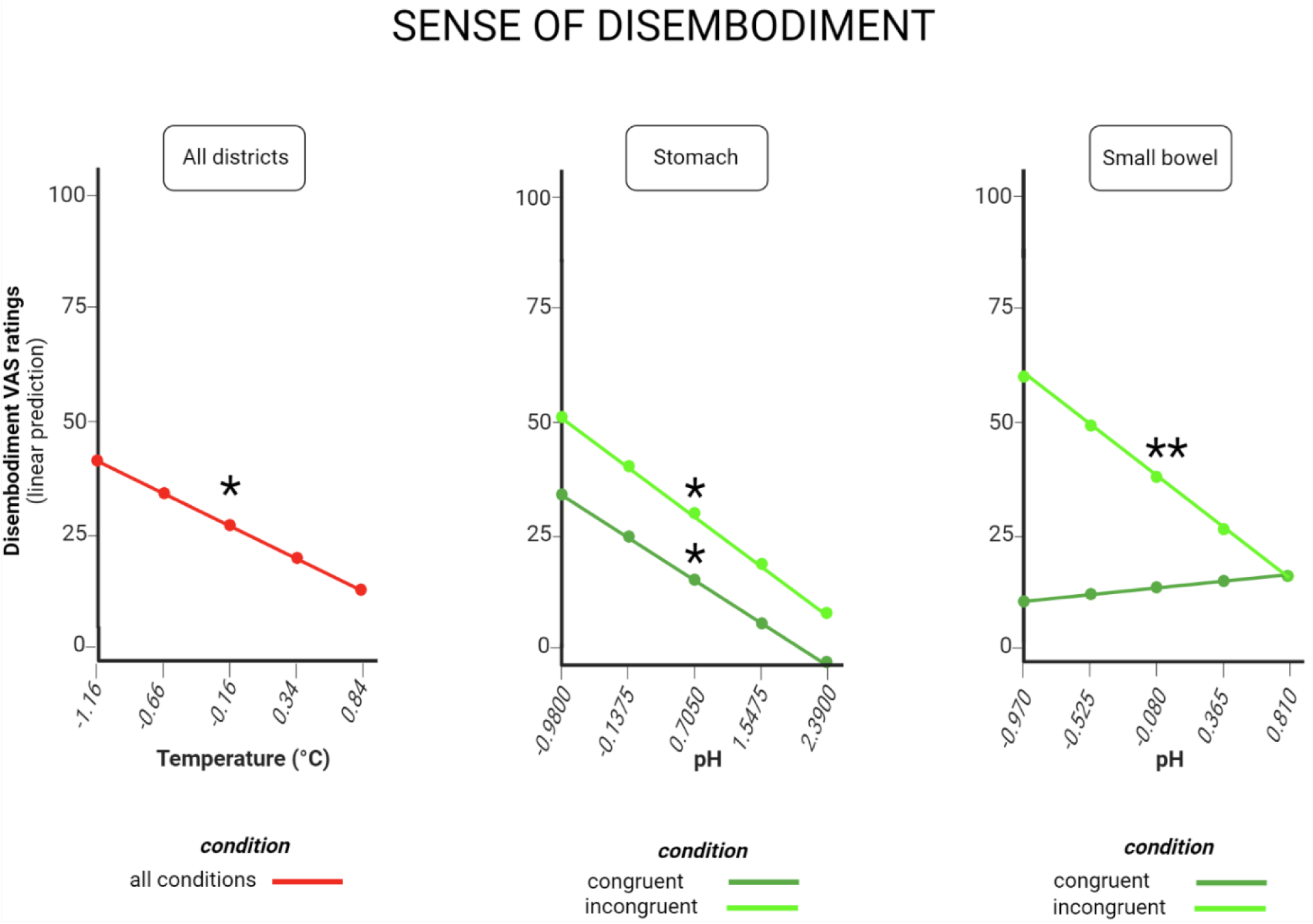
Effects of temperature (left), stomach pH (centre) and small bowel pH (right) on feelings of disembodiment, as assessed through Visual Analogue Scales (VAS) ratings (estimated marginal means). ‘All districts’ indicates that the effect holds true across all the three regions of the gastro-intestinal tract we tested (stomach, small bowel, and large bowel). ‘Congruent condition’ means that the effect shown in light green occurs when participants observe a congruent virtual body that matches their physical body. ‘Incongruent condition’ means that the effect shown in dark green occurs when participants observe an incongruent virtual body that does not match their physical body. ‘All conditions’ means that the effect occurs both for congruent and incongruent condition. * p < .05 ** p <.01

Furthermore, participants felt less disembodied also when the pH of their stomach or small intestine became less acidic. For each 1-unit increase in stomach pH, predicted VAS ratings of disembodiment fell by 11.39 ± 5.36 points when the virtual body matched the physical body of the participant (congruent condition, *t* = −2.124, *p* = 0.0340; **Fig. 2** centre and **Table S4**) and by 13.20 ± 6.32 points when it did not (incongruent condition, *t* = −2.089, *p* = 0.0370; **Fig. 2** centre and **Table S4**). Likewise, for each 1-unit decrease in small bowel pH, predicted VAS ratings of disembodiment dropped by 25.00 ± 9.52 points in the incongruent condition (*t* = −2.627, *p* = 0.0088; **Fig. 2** right and **Table S4**).

#### Sense of location covaries with gut pressure and pH

The higher the pressure across the gastro-intestinal tract, the more our participants felt they occupied the same place of the virtual body. For each 1 mmHg increase in stomach, small bowel, or large bowel pressure, predicted VAS ratings of the feeling of being located into the virtual body went up by 5.62 ± 2.25 points (*t* = 2.496, *p* = 0.0128; **Fig. 3** left and **Table S3**).

**Figure 3.**
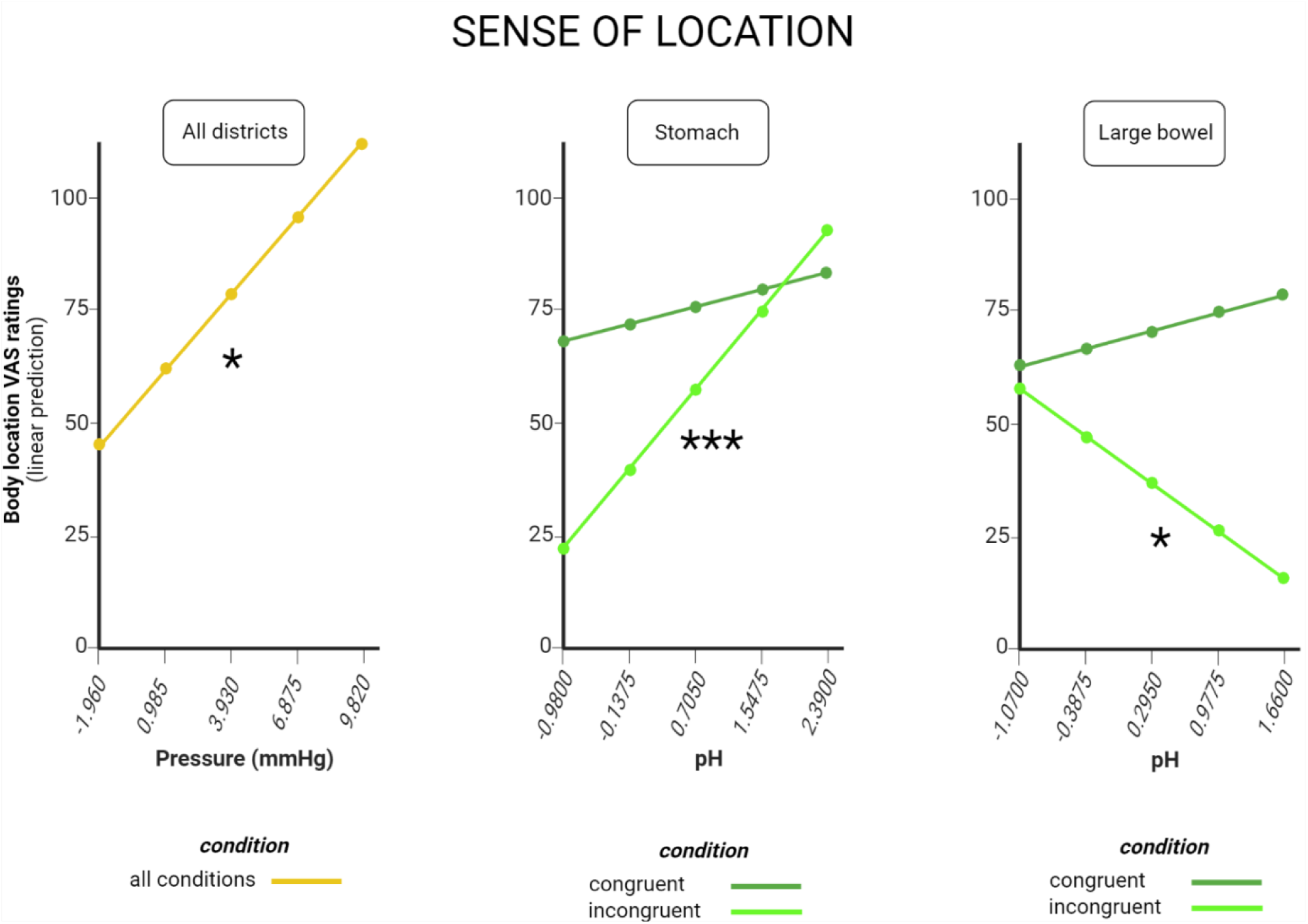
Effects of pressure (left), stomach pH (centre) and large bowel pH (right) on feelings of location, as assessed through Visual Analogue Scales (VAS) ratings (estimated marginal means). ‘All districts’, ‘congruent condition’, ‘incongruent condition’, and ‘all conditions’ defined as in Figure 2. * p < .05 *** p <.001

A stronger sense of location correlated also with less acidic pH of the stomach and more acidic pH of the large bowel when the virtual body did not match the real body (incongruent condition). For each 1-unit increase in stomach pH, predicted VAS ratings of being located in the incongruent virtual body increased by 21.02 ± 6.32 points (*t* = 3.326, *p* < 0.001; **Fig. 3** centre and **Table S4**). For each 1-unit increase in large bowel pH, the same ratings went down by 15.44 ± 6.56 points (*t* = −2.355, *p* = 0.0188; **Fig. 3** right and **Table S4**).

#### Sense of agency covaries with small bowel pH

The less acidic the small bowel pH, the more our participants felt they were in control of the incongruent virtual body. In the incongruent condition, for each 1-unit increase in small bowel pH, predicted VAS ratings of agency rose by 24.21 ± 9.52 points (*t* = 2.545, *p* = 0.0111; **Fig. 4** left and **Table S4**).

**Figure 4.**
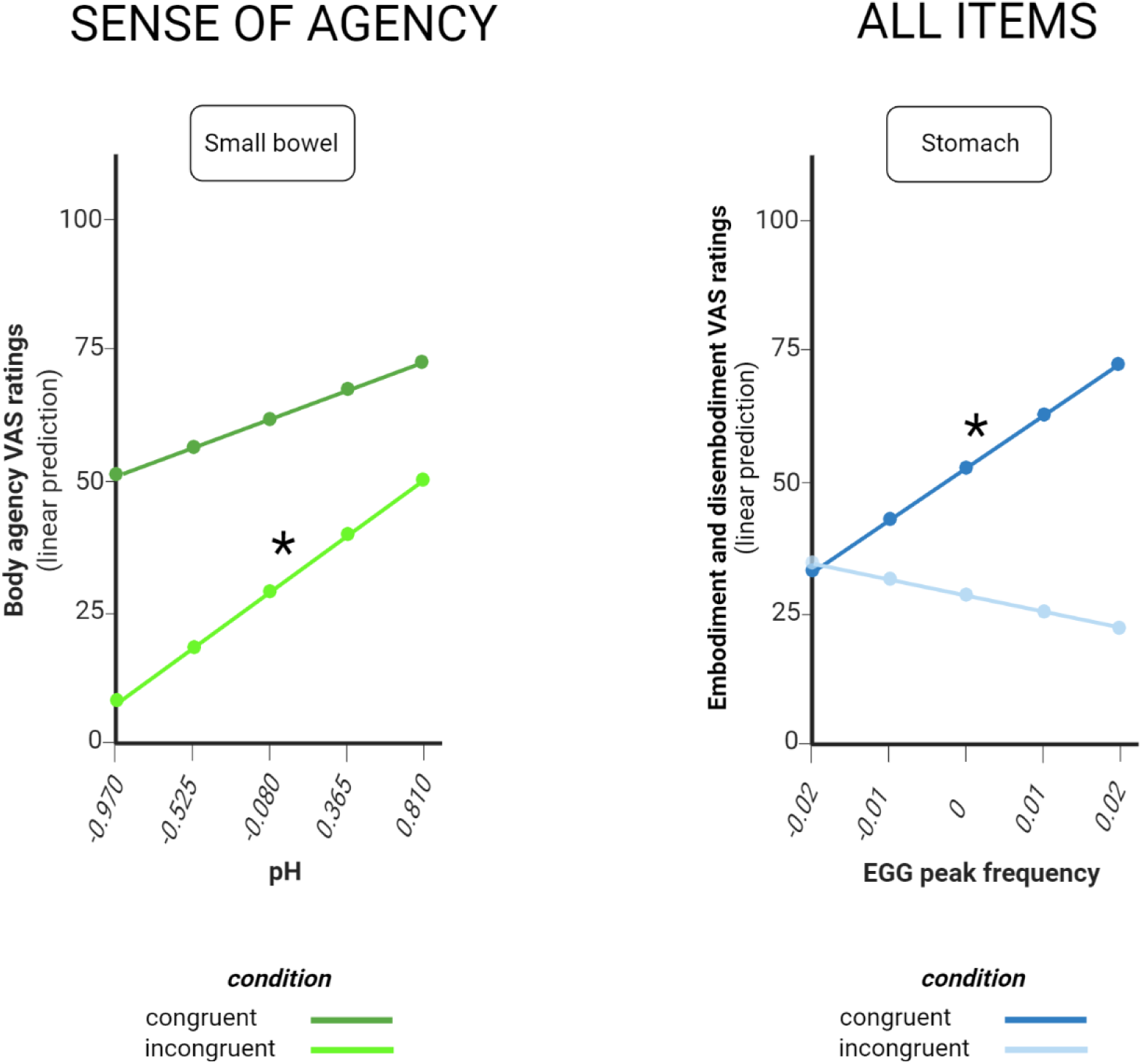
Effect of small bowel pH on agency ratings (left) and effect of EGG peak frequency on all embodiment and disembodiment ratings (right). ‘Congruent’ and ‘incongruent’ condition defined as in figure 2. * p < .05

### Model 2 (EGG) results

#### All items of the BSC questionnaire covary with stomach peak frequency

When participants embodied a virtual body matching their real body, the higher the peak frequencies of gastric myoelectric activity, as recorded through electrogastrography (EGG), the higher the ratings across all items of the bodily self-consciousness (BSC) questionnaire. For each 0.01 Hz increase in the frequency of gastric contractions, predicted VAS ratings across the BSC questionnaire increased by 9.72 ± 3.84 points (*t* = 2.534, *p* = 0.0122; **Fig. 4** right and **Table S6**).

## Discussion

Capitalizing on the smart pills technology to wirelessly monitor the physiological parameters of the gastro-intestinal tract in real time while healthy subjects underwent a breathing-related bodily illusion in immersive virtual reality, we were able to find an association between individual ratings of bodily self-consciousness and individual patterns of gut activity. According to our results, physiological parameters of the stomach and the bowel were positively associated to the perception of embodiment and showed the opposite pattern in relation to the feeling of disembodiment. Specifically, when gastro-intestinal pressure and gastric pH increased, so did the sense of location. When small bowel pH rose, the sense of agency rose too. Instead, when pH and temperature went up, participants reported to feel less disembodied.

Higher intraluminal pressure and temperature indicate that the gastro-intestinal tract is more active. In particular, higher pressure is a sign that gut contractions have become stronger and more frequent^12^, while higher temperature is associated with an increase in metabolic rate^22^. Thus, stomach and bowel activity are directly correlated with the feeling of location and inversely correlated with the feeling of disembodiment. The fact that gut activity scales with some facets of bodily self-consciousness is consistent with analogous trends observed in other physiological domains, including respiration^9^, histamine reactivity^23^, midterm skin temperature^24^ and skin conductance^25–27^. Signals relayed from more active organs and tissues may boost the salience and strength of body representations in target cortical areas. In turn, reinforced body representations may reduce the feeling of having no body and increase the sensation of dwelling into a body.

While temperature and pressure are gut-wide markers of bodily self-consciousness, displaying the same correlation with location and disembodiment across organs and experimental conditions, the role of pH tends to be selective for the incongruent condition and changes from organ to organ. When participants observe an incongruent virtual body, higher (i.e., less acid) stomach pH is correlated with higher location and lower disembodiment; higher small bowel pH goes hand in hand with agency; and higher large bowel pH is inversely correlated with location.

Of note, pH values have different meanings in different organs. In particular, higher pH indicates *less* activity in the stomach, since gastric acid secretion, a hallmark of an active stomach^28^, would entail a lower pH. Less gastric activity may temporarily induce a looser representation of the real body, thus allowing participants to feel localised in an incongruent virtual body and reducing their disembodiment when they observe the incongruent avatar. On the contrary, higher pH could indicate *more* activity in the bowel, whose processes take place in a basic environment^29^. More bowel activity may sharpen the representation of the real body and thus make participants less likely to feel localised in an incongruent virtual body, while also reducing disembodiment. However, it is unclear why an increase of small bowel activity should lead to increased agency. More generally, it may also be that physiological activation is an effect, rather than a cause, of higher embodiment and lower disembodiment. In fact, embodiment and physiology may be engaged in a positive feedback cycle. New experimental protocols that stimulate or inhibit the activity of the stomach and the bowel (e.g. presenting pictures^30^, evoking food-related thoughts^31^ or electrically stimulating the vagal afferents^32,33^) are needed to shed light on causality.

Whilst physiological signals recorded by the SmartPill were tied to specific facets of bodily self-consciousness, a surge in stomach contractions, as measured through EGG peak frequency, led to a broad, generalised increase of both embodiment and disembodiment ratings. Thus, compared to EGG, smart pills offer a much more nuanced picture of the links between gut physiology and the bodily self. In principle, the same approach could be extended beyond bodily self-consciousness, harnessing the power of ingestible capsules to gauge the correlation between gut states and mental states with unprecedented detail. Hence, together with other ingestibles^34–38^, smart pills could become a particularly valuable, minimally invasive tool for assessing also other important psychological constructs that may be rooted in gastro-intestinal physiology, from emotions^39^ to moral disgust^40,41^.

Overall, our findings indicate that physiological information coming from the whole gastro-intestinal tract is deeply linked to one’s sense of oneself as an embodied entity. This suggests that bodily self-consciousness is based upon multisensory integration processes whose scope is even wider than previously thought – processes gathering a wealth of sensory information from abdominal viscera as well as from thoracic organs and from skin receptors. Such a diverse range of sensory inputs probably helps the brain to maintain a ‘body template’ in a variety of circumstances, including those in which a particular sensory channel may be temporarily or permanently shut down, as in darkness, blindness, or spinal cord injuries. In these cases, spared connections between the gastro-intestinal tract and the cortex may preserve a correct representation of the body despite the lack of exteroceptive sensory information. By the same token, this compensatory process may be impaired in psychiatric conditions that alter bodily self-consciousness, as in eating disorders and depersonalization. Applying the current experimental paradigm to clinical samples will enable researchers to ascertain the impact of impaired gut-brain signalling on corporeal awareness and lay the groundwork for future rehabilitation protocols.

Given the sex differences in the perception of signals coming from internal organs^42,43^ and in EGG frequency^44^, recruiting only male participants allowed a more stringent test of the role of gastro-intestinal signals in bodily self-consciousness. Nevertheless, we are aware that future experiments should involve a larger sample of participants of both sexes to increase the generalizability of the present findings. Finally, although the ingestible device is minimally invasive, its dimensions and sampling rate are sub-optimal. Ongoing advances in pill technology may overcome this issue and deepen our understanding of the connection between our gut and our bodily self.

## Methods

### Experimental design

#### Participants

Thirty-one healthy male subjects aged 20-30 years (M = 24.42, SD = 2.8) took part in the study after giving their informed consent. No subject had a history of psychiatric or neurologic disorders. A detailed screening procedure ensured that all participants were eligible for ingesting the SmartPill™ without any known contraindication (history of gastric bezoars; history of any abdominal/pelvic surgery within the previous three months; swallowing disorders; suspected or known strictures, fistulas, or physiological/mechanical obstruction within the gastro-intestinal tract; dysphagia to food or pills; Crohn’s disease or diverticulitis; body mass index ≥ 40; and cardiac pacemakers or defibrillators^12^). All subjects were naïve to the purpose of the research and were paid for their participation. The study was approved by the local Institutional Review Board (Fondazione Santa Lucia ethics committee).

#### Materials

##### 1. SmartPill™

The gastro-intestinal milieu of each participant was monitored through a SmartPill™ (SmartPill Motility Testing System, Medtronic plc). SmartPills are light, single-use, orally ingestible capsules (length: 26 mm; width: 13 mm; weight: 4.5 g). Each pill consists of a polyurethane shell fitted with a long-lasting battery (> 5 days), a transmitter (broadcast frequency: 434.2 MHz), and internal sensors probing temperature (range: 20-42 °C; accuracy: ± 1 °C), intraluminal pressure (range: 0-350 mmHg; accuracy: ± 5 mmHg in the 0-99 mmHg sub-range, ± 10% of applied pressure in the 100-350 mmHg sub-range) and pH (range: 1-9; accuracy: ± 0.5 pH units) of the gastro-intestinal (GI) tract (**Fig. 1B** and **Fig. 5A**).

**Figure 5.**
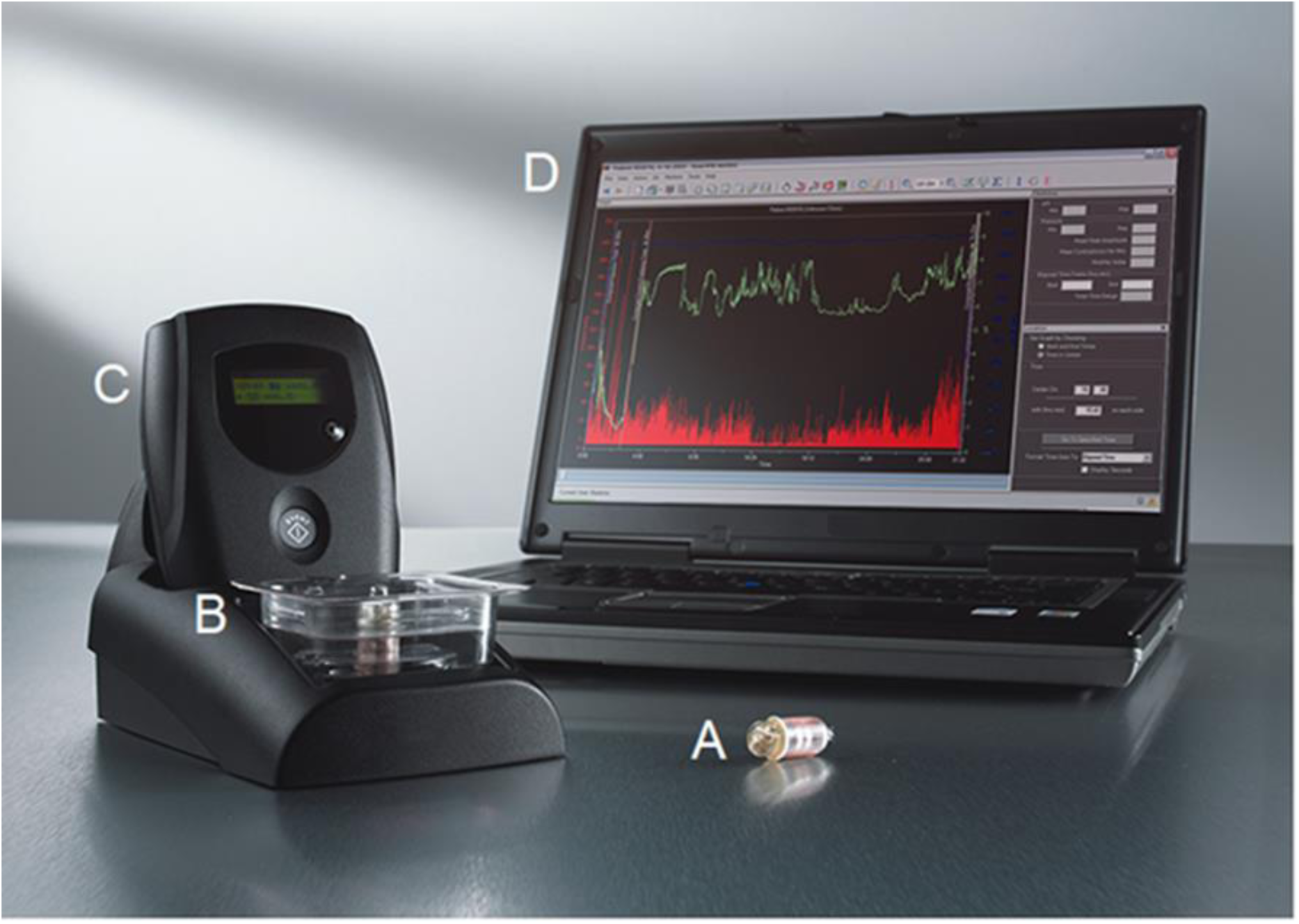
SmartPill apparatus. A: capsule. B: pH calibration buffer solution. C: data receiver and logger. D: laptop with graph showing pH (green line), temperature (blue line) and pressure (red bars) over time. Note the abrupt rise of the pH values in the left part of the screen, signalling the passage from the stomach to the small bowel, the slow build-up of the signal in the small bowel, and the rapid decrease marking the entrance of the pill in the large bowel. Adapted from a picture of Medtronic plc.

Before being ingested, the pill is activated through a magnetic fixture and the pH sensor is calibrated through a buffer solution (**Fig. 5B**). After ingestion, the capsule samples temperature data every 20 s, pressure every 0.5 s, and pH every 5 s for the first 24 hours; sampling frequencies are halved thereafter. The pill transmitter wirelessly sends these data to an external radio receiver (operating range: ~ 1.5 m), which can be either docked in a dedicated station or comfortably fastened to a belt worn by the participant (**Fig. 5C**).

Combining pH, pressure, and temperature information, the MotiliGI software (Medtronic plc) univocally identifies the specific segment of the GI tract in which the pill is located at a given time. The software takes an abrupt increase of ≥ 2 pH units as a sign that the pill left the stomach and entered the small intestine. Likewise, the software interprets a subsequent gradual decrease of ≥ 1 pH unit for at least 10 consecutive minutes as a sign that the pill left the small intestine and entered the large intestine (**Fig. 5D**).

In our sample, 30 out of 31 subjects displayed a pH increase and decrease as expected, while for the remaining subject the software was still able to localize the GI districts that the pill went through based on pressure, temperature, and time data.

##### 2. Electrogastrography (EGG)

Electrogastrographic (EGG) recordings were used as a measure of gastric contractions. EGG records the electrophysiological activity of a selected cluster of cells at the junction of the enteric nervous system with the stomach – the so-called interstitial cells of Cajal (ICC). ICC act as pacemakers of stomach contractions by generating and propagating electric slow waves that have a normal frequency of 0.05 Hz, i.e. 3 cycles/minute^45,46^. In healthy subjects, each slow wave is coupled to a gastric contraction^21^. Slow wave electrical signals were recorded through a standard 1-channel EGG bipolar montage^21^ with 3 pre-gelled disposable Ag/AgCl electrodes. Participants were instructed to lie supine on a deck chair, then their abdominal skin was accurately cleansed to reduce impedance. The first recording electrode was placed halfway between their xyphoid and their umbilicus, while the second recording electrode lay 5 cm up and 5 cm to the left of the first (taking the left side of participants as a reference) and the ground electrode lay on the left costal margin (**Fig. 6**).

**Figure 6.**
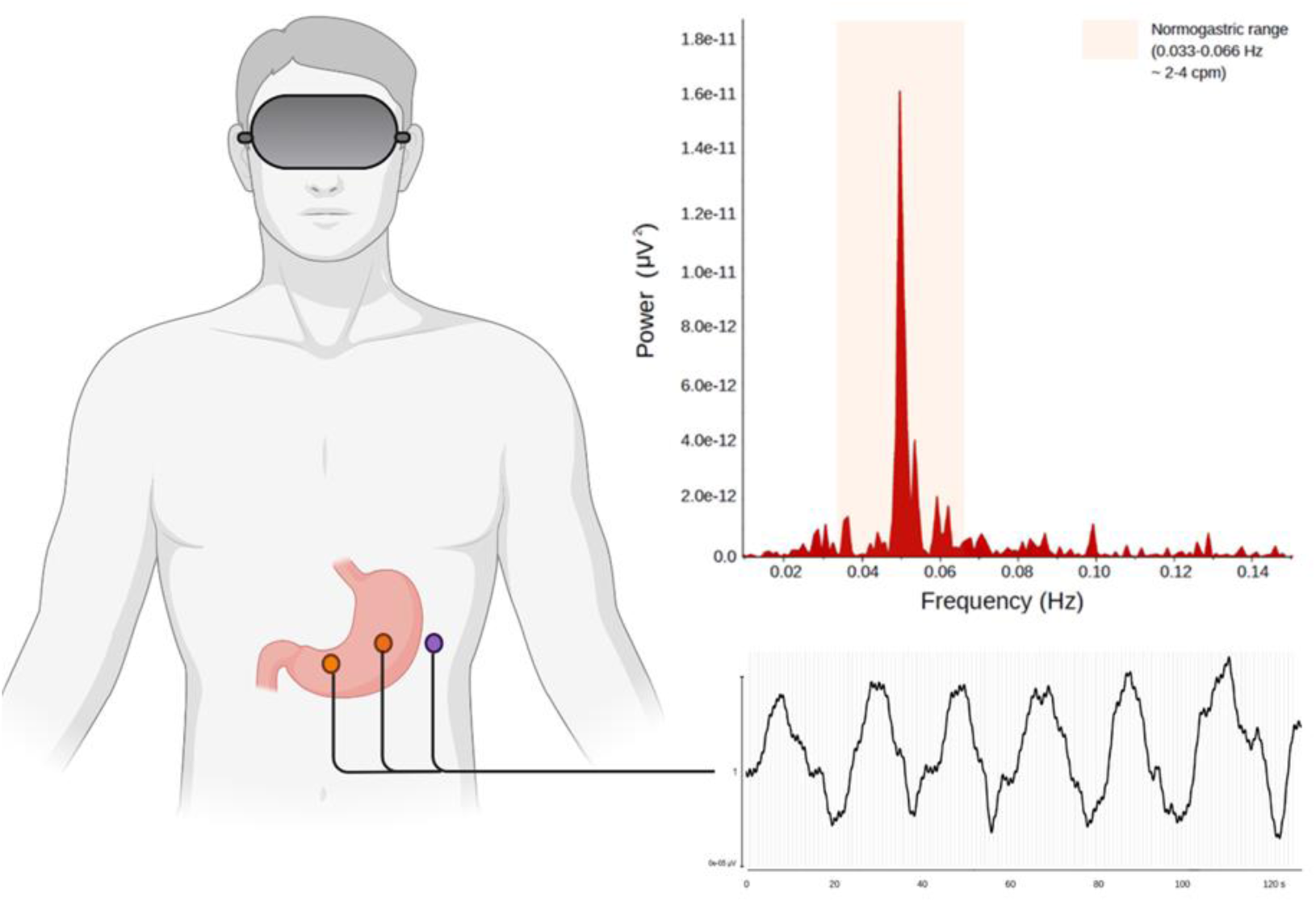
1-channel EGG bipolar montage (left) with two recording electrodes (orange dots) and one ground electrode (purple dot), sample EGG recording (bottom right), and sample EGG periodogram (top right). Samples collected from a participant in our lab.

##### 3. Immersive virtual reality

The immersive virtual reality apparatus for the embreathment illusion^9^ included a virtual scenario designed in 3DS Max 2015 (Autodesk Inc) and Unity 2017.1 (Unity Technologies SF). The scenario was broadcasted to a VIVE headset (HTC Corp., 6 degrees of freedom, field of view: ~110°, resolution: 2160×1200 (1080×1200 per eye, aspect ratio 9:5), refresh rate: 90 Hz) and consisted of a life-size room in which a virtual body (avatar) lay on a deck chair.

In the *congruent* condition, the avatar was seen from a first-person perspective, had a human-like appearance, and breathed as the participant, i.e. it inspired when the participant inspired and expired when the participant expired, in real time. The exact alignment of real and virtual breathing was obtained through a customized VIVE sensor (**Movie S1**) that mapped real, respiration-induced belly movements onto the virtual body with sub-millimetric precision (error < 10^−3^ m).

In the *incongruent* condition, the avatar was seen from a third-person perspective, had an object-like wooden appearance, and breathed in anti-phase with the participant, i.e. it expired when the participant inspired and vice versa, in real time. The same VIVE sensor described above was adopted also in this condition, although in this case the y-axis of the sensor was mapped in a reverse fashion onto the virtual body to further enhance the incongruity effect.

A custom graphical user interface (GUI) was embedded in the virtual scenario, allowing participants to answer some questions relative to bodily self-consciousness (see section 4 below) at the end of each condition. For a detailed footage of the immersive virtual reality experience, please see **Movie S1**.

##### 4. Measures of bodily self-consciousness

At the end of each virtual reality condition, we administered a customized bodily self-consciousness questionnaire^9^ consisting of five different 0-100 visual analogue scales (VAS). In each scale, participants indicated how much they agreed with a statement by selecting a point on a line ranging from complete disagreement (leftmost point) to complete agreement (rightmost point) through a joystick. **Table 1** shows the complete list of statements.

##### 5. Measures of interoception

We also measured objective^47^ and subjective^48^ levels of interoception, i.e. how participants perceived their inner physiological signals^49^ (see **Supplementary Information** for details on methods and results and **Table S7** for descriptive statistics).

#### Data collection procedure

To ensure that the data gathered by the SmartPill were as reliable as possible, subjects were instructed to discontinue any medication that could interfere with pH values and gastro-intestinal motility^12^. Specifically, we checked that none of the participants was assuming any: i) proton pump inhibitors in the seven days before the experiment; ii) antihistamines, prokinetics, antiemetics, anticholinergics, antidiarrheals, narcotic analgesics, and non-steroidal anti-inflammatory drugs in the three days before the experiment; iii) laxatives in the two days before the experiment. Participants were instructed not to take antacids and any alcohol the day before the experiment. Eight hours before the experiment, they also stopped eating and smoking.

The day of the experiment, participants came to the laboratory, filled in the informed consent form, and ate a standardized ~260 kcal breakfast consisting of egg whites (120 g), two slices of bread, and jam (30 g) to make sure that gastro-intestinal transit times of the SmartPill were not affected by meal variability. Meanwhile, we activated the capsule through a magnetic fixture and calibrated the capsule pH sensor (see Materials above, section 1).

After calibration was complete, the pill started transmitting data to the radio receiver. Data came with a relative time stamp indicating the number of seconds elapsed from calibration, but no absolute time reference. To overcome this issue, we synchronized calibration with an external clock that provided us with the required absolute time frame. At that point, participants swallowed the SmartPill while drinking a glass of water (120 ml). A medical doctor supervised the ingestion procedure to help in case of swallowing problems. All subjects ingested the pill without any trouble. After the ingestion, participants fastened the receiver around their belt and lay supine on a deck chair. This allowed the experimenters to place EGG electrodes according to the montage described above (see Materials, section 2). When the whole apparatus was in place, we recorded a 15-minute resting-state SmartPill/EGG baseline session in which participants were instructed to relax and keep their eyes open. Then, we perused real-time pH data displayed on the receiver to make sure that the capsule was working and actually lay in the stomach, as signalled by a highly acidic pH (~1-2).

After this requirement was fulfilled, we administered a simplified version of the embreathment illusion^9^ delivered through a virtual reality headset and a customized breathing sensor (see Materials, section 3). Both the *congruent* and the *incongruent* condition of the illusion (see above) lasted for 240 s and were followed by the bodily self-consciousness questionnaire described in section 4 of the Materials. The order of experimental conditions was counterbalanced across participants. Conditions were interspersed with 5’ washout pauses to avoid carryover effects. Throughout each experimental condition, the receiver logged SmartPill data about the pressure, temperature, and pH of the stomach, while a dedicated amplifier (ADInstruments PowerLab) registered the EGG signal (**Fig. 7**).

**Figure 7.**
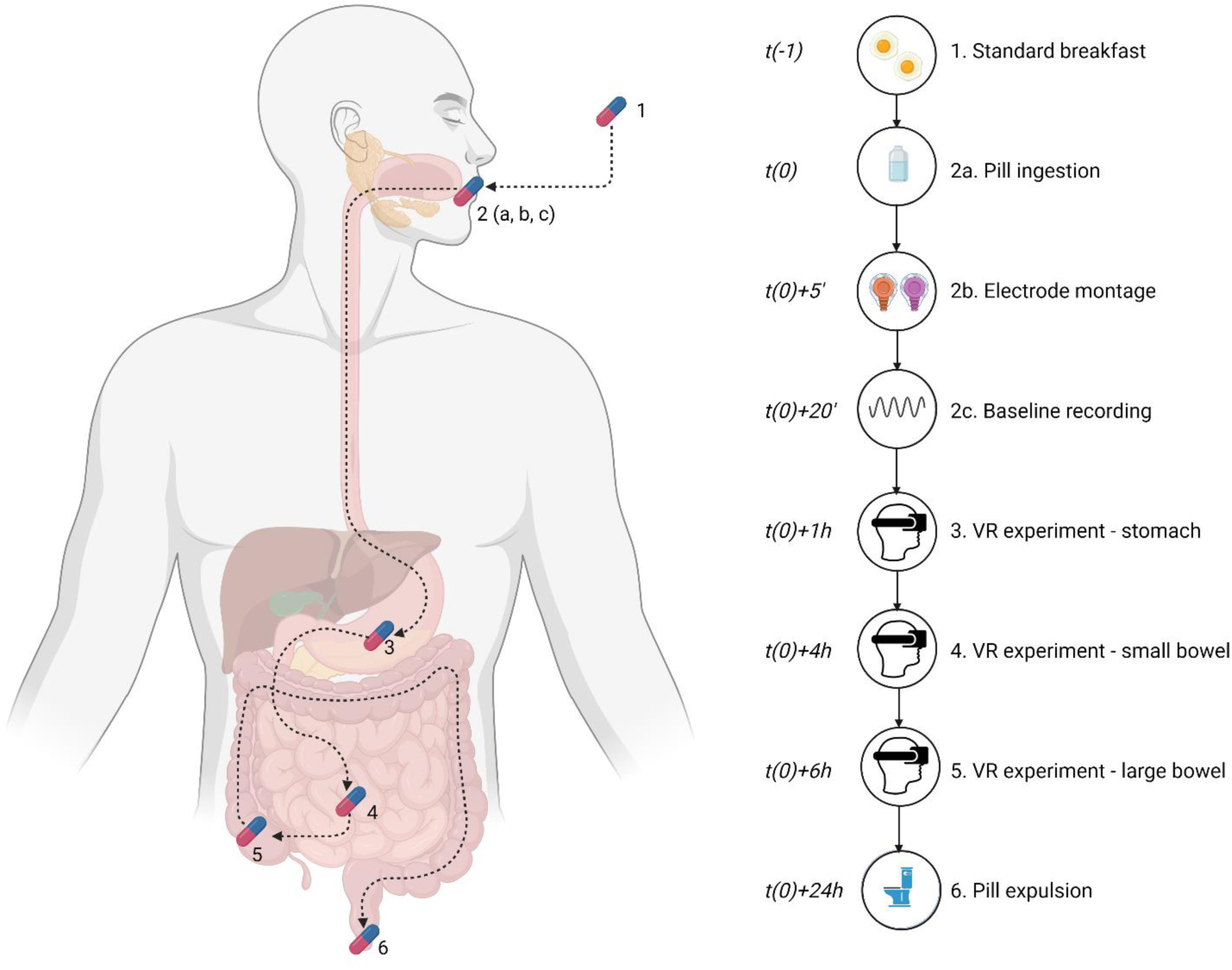
Timeline of the experimental procedures. Italicized text indicates the approximate average time at which each event occurs.

As we were interested in assessing the coupling between bodily self-consciousness and the physiology of each main segment of the gastro-intestinal tract, we waited until the pill went through the pylorus (as marked by a ≥ 2 pH units sudden increase: see above) to repeat the virtual reality experience. This normally occurred within 2-5 hours from the ingestion of the capsule. At that point, we administered again the embreathment illusion, this time recording only small bowel data from the SmartPill. When the capsule entered the large bowel (as marked by a ≥ 1 pH unit decrease lasting for at least 10 minutes, typically observed after 2-6 hours from the stomach-small bowel transition: cf. above), we administered the illusion again for the third and last time, always logging SmartPill data for each experimental condition (**Fig. 7**).

After the first 6 hours from the beginning of the experiment, participants were provided with a meal. After the first 8 hours, they could smoke again. After 3 days, they were allowed to drink alcohol as usual. During the pauses between the stomach and small bowel data collection and between the small and large bowel, they underwent subjective and objective tests of interoception (see Materials, section 5 and Supplementary Information) and then were free to work or study as they pleased, although they had to avoid strenuous physical exercise. Finally, after the last experimental condition of the large bowel was over, participants could leave the lab. However, they kept the receiver with them, so that they could check the gut physiological parameters for themselves until the capsule stopped transmitting data and was expelled through defecation, ordinarily 10-73 h after ingestion^11^.

### Data analysis procedure

#### SmartPill data pre-processing

Raw SmartPill data were downloaded from the receiver and exported as .txt files. A custom MATLAB algorithm converted relative timestamps in absolute times, so that each event (e.g. beginning and end of each experimental condition) was paired to a definite hh:mm:ss:ms string. We computed the gastric, small bowel, large bowel, and whole gut transit times of the capsule^11^ to check whether any subject displayed anomalies in their gastric physiology. 30 out of 31 subjects had normal transit times, while the remaining subject had an abnormal large bowel transit time (>> 59 h, cf. ^12^). Consequently, his SmartPill data were discarded.

#### EGG data pre-processing

Raw EGG recordings were visually inspected to remove artifacts due to body movements. A 0.016-0.15 Hz bandpass filter removed pink noise and unwanted higher frequencies that are ordinarily associated with cardiac, respiratory, and small bowel activity (cf. ^50^). The artifact-free tracings thus obtained were then used to extract the EGG peak frequency for each subject and experimental condition (**Fig. 6**). EGG spectral density was computed using Welch’s method on 200 s time windows with 150 s overlap^18^. EGG peak frequency was defined as the maximum periodogram peak in the ‘normogastric’ range, i.e. the range of frequencies that is compatible with the number of stomach contractions in healthy individuals (0.033– 0.066 Hz ~ 2-4 cycles per minute; cf. ^18^). The whole EGG analysis procedure was performed with BrainVision Analyzer (Brain Products GmbH) and the MATLAB FieldTrip toolbox^51^.

#### VAS questionnaire data pre-processing

Raw bodily self-consciousness ratings provided by the subjects through the VAS questionnaire in the virtual reality GUI were exported, then matched with the average pH, pressure, temperature, and EGG peak frequency values computed for each participant, each experimental condition, and each gastro-intestinal district (stomach, small bowel, large bowel). The resulting data matrices are available at https://osf.io/wecta/?view_only=45e1a9e30c2a47efa82c10161b70b732

#### Statistical data analysis

We used R (version 3.6.1) and the R *lme4* package^52^ to perform a linear mixed-effects analysis of the data. We modelled how much ratings of perceived body ownership, agency, location, disembodiment, and multiple bodies (see above and **Table 1**) changed depending on the experimental conditions, the gastro-intestinal (GI) region, and, most importantly, the mean pH, pressure, temperature, and peak frequency values recorded in each experimental condition.

We built two distinct mixed-effects models. Model 1 (SmartPill model) tested the influence of pH, pressure, and temperature of the three GI regions (stomach, small bowel, and large bowel) over bodily self-consciousness. The dependent variable was the whole set of bodily self-consciousness VAS ratings. As fixed effects, the model had the experimental *condition*, i.e. human-avatar sensory congruency (two levels: congruent and incongruent), the VAS *item* (five levels: perceived ownership, agency, location, disembodiment, and two bodies: see **Table 1**), the GI region (three levels: stomach, small bowel, large bowel) and the condition-specific *pH* (continuous), *pressure* (continuous), and *temperature* (continuous).

Model 2 (EGG model) assessed the influence of EGG peak frequencies over bodily self-consciousness. Since EGG peak frequencies are a specific measure of stomach activity, here the dependent variable was a subset of bodily self-consciousness VAS ratings, namely, those collected after the first embreathment illusion, when the pill was in the stomach. Like Model 1, also Model 2 featured condition, item, and GI region as factors, but replaced pH, pressure, and temperature with condition-specific *EGG peak frequencies*.

In both models, fixed effects were tested for interactions with each other. Continuous fixed effects were also group-mean-centred, subtracting the mean of stomach pH from stomach pH values, the mean of small bowel pH from small bowel pH values, and so forth. As random effects, both models included by-subject random intercepts. Model 1 also had by-condition random slopes. Overall, each mixed model was specified as follows.

Model 1 (SmartPill)

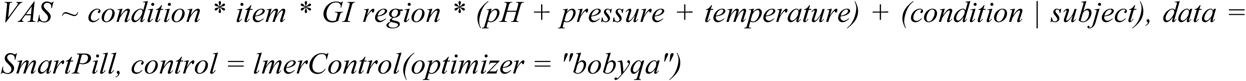

Model 2 (EGG)

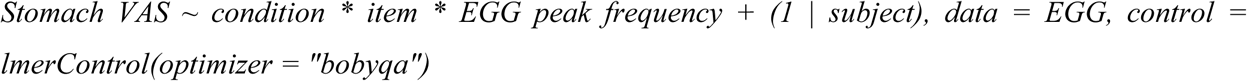

To minimise the chance of Type 1 errors^53^, models were fitted with Restricted Maximum Likelihood (REML) and *p* values were obtained through Type II analysis of variance with Satterthwaite’s method^54^, as implemented in the *lmerTest* package^55^. Statistically significant interactions were followed up with post-hoc tests of the simple effects involving gut physiological parameters, experimental conditions, and bodily self-consciousness ratings against the null hypothesis of a slope equal to zero. All post-hoc tests used Kenward-Roger degrees of freedom^56^ and were performed via the *emmeans* package to obtain estimated marginal means (EMMs). EMMs were then plotted with the *emmip* function.

Both Model 1 and Model 2 converged successfully. The standard assumptions and requirements of mixed models (linearity, homoscedasticity, absence of collinearity, and normality of residuals) were assessed through visual inspection of residual plots, the *shapiro*.*test* function, and the *vif* function. The percentage of variance explained by each mixed-effects model^57^ was computed through the *r*.*squaredGLMM* function of Kamil Bartoń’s MuMIn Package.

## Data availability statement

The data that support the findings of this study are available in Open Science Framework (OSF), https://osf.io/wecta/?view_only=45e1a9e30c2a47efa82c10161b70b732

## Author Contributions

Conceptualization and methodology: A.M., G.P., M.S.P., S.M.A. Data curation, formal analysis, investigation: A.M., G.P., M.S.P. Funding acquisition, project administration, resources, supervision: SMA. Software, visualization, writing – original draft: A.M. Writing – review & editing: A.M., G.P., M.S.P., S.M.A.

## Acknowledgments

We thank Danila Cosenza for medical assistance and Maurizio Molisso for standardized meal preparation. Figures designed with BioRender.com. Supported by ERC Advanced Grant 789058 (to S.M.A.).

