## Supplemental Information for "Gut markers of bodily self-consciousness"

### Supplementary Data

Please find below an extended technical description of the results presented in the main text.

#### 1. SmartPill results

Model 1 (cf. Methods section of the main manuscript) converged successfully with no errors and no warnings. The model had a marginal $R^2$ = .40 and a conditional $R^2$ = .61. Visual inspection of the residual plots did not reveal any obvious deviation from homoscedasticity. Residuals were not normally distributed (Shapiro-Wilk normality test, $p < .05$), but linear models are robust against violations of normality[1]. As for collinearity, all independent variables and their products had a GVIF^(1/(2*Df)) < 7. The ratio between the number of data points ($n$ = 855) and the number of estimated parameters ($k$ = 120) was $n/k$ = 7.125, above the minimum ratio of $n/k$ = 3 (see [2]). Thus, the model was not overparametrized.

Type II analysis of variance of Model 1 yielded statistically significant 2-way interactions between *item* and *pressure* ($F = 3.0497$, $p = 0.0166$) and between *item* and *temperature* ($F = 3.0218$, $p = 0.0174$). Furthermore, there was a statistically significant 4-way interaction between *condition*, *item*, *gastro-intestinal region*, and *pH* ($F = 2.4228$, $p = 0.0139$) (**Table S1**). The post-hoc tests of these statistically significant 2-way and 4-way interactions, whose key results are already described in the main text, are presented in **Tables S2-S4**.

Other significant 3-way and 2-way interactions, as well as significant main effects, are listed in **Table S1** and were not further discussed due to the presence of the significant higher-order (2-way or 4-way) interactions described above.

### 2. Electrogastrography (EGG) results

Model 2 (cf. Methods section of the main manuscript) converged successfully with no errors and no convergence warnings. The ratio between the number of data points ($n = 260$) and the number of estimated parameters ($k = 20$) was $n/k = 13$, above the minimum ratio of $n/k = 3$ (see [2]). Thus, the model was not overparametrized.

Type II analysis of variance of Model 2 yielded statistically significant 2-way interactions between *condition* and *item* ($F = 15.8484$, $p < 0.001$) and between *condition* and *EGG peak frequency* ($F = 4.6064$, $p = 0.033$) (**Table S5**). Other significant effects are listed in Table S5 and were not further discussed due to the presence of significant higher-order interactions. Post-hoc tests of the c*ondition x EGG peak frequency* interaction, whose key result is already mentioned in the main text, are presented in **Table S6**.

### 3. Interoception results

Participants displayed ordinary levels of interoception (**Table S7**). In particular, Shapiro-Wilk tests indicated that interoceptive sensibility, accuracy, and awareness scores did not significantly deviate from a normal distribution (interoceptive sensibility: $W = 0.969$, $p = 0.565$; interoceptive accuracy: $W = 0.941$, $p = 0.086$; interoceptive awareness: $W = 0.978$, $p = 0.753$).

### Supplementary Methods

*Interoceptive sensibility*, that is, the participants' subjective, self-reported ability to monitor interoceptive signals[42], was measured through the Italian version of the Multidimensional Assessment of Interoceptive Awareness (MAIA)[22], a list of 32 questions sampling how much each participant is aware of their physiological state. In particular, the average of three MAIA subscales (*Noticing*, *Attention Regulation*, and *Body Listening*) was taken as a proxy of interoceptive sensibility. Responses were provided with a 6-points Likert scale.

*Interoceptive accuracy*, that is, the participants' objective performance at perceiving interoceptive signals[42], was assessed via Schandry's heartbeat counting task[21]. Subjects were asked to report the number of heartbeats they perceived in four different time windows (25 s, 35 s, 45 s, 100 s) without guessing or relying on external cues (e.g. taking their own pulse). Meanwhile, EGG electrodes were used to pick objective electrocardiographic (EKG) signals. Raw EKG recordings were processed in LabChart to detect the QRS complex associated with each heartbeat and thus compute the actual number of heartbeats for each time window. These objective data were then paired with self-reported numbers of heartbeats for each time window to calculate an interoceptive accuracy score for each participant ranging from 0 (not accurate at all) to 1 (perfect accuracy).

*Interoceptive awareness*, that is, the participants' confidence in how well they fared in the heartbeat counting task, was gauged through a visual analogue scale (VAS) that ranged from 0 (indicating participants believed their performance was extremely poor) to 100 (indicating participants believed their performance was extremely good).

| | *Sum Sq* | *Mean Sq* | *NumDF* | *DenDF* | *F* | *p* |
|---|---|---|---|---|---|---|
| condition | 30123 | 30123 | 1 | 26.58 | 69.4735 | < 0.001 *** |
| item | 138897 | 34724 | 4 | 682.05 | 80.0860 | < 0.001 *** |
| region | 2500 | 1250 | 2 | 689.12 | 2.8825 | 0.0566674 |
| ph | 12 | 12 | 1 | 720.92 | 0.0271 | 0.8693748 |
| pressure | 176 | 176 | 1 | 732.28 | 0.4055 | 0.5244588 |
| temperature | 233 | 233 | 1 | 178.07 | 0.5365 | 0.4648315 |
| condition:item | 104086 | 26022 | 4 | 682.05 | 60.0145 | < 0.001 *** |
| condition:region | 440 | 220 | 2 | 689.35 | 0.5079 | 0.6019753 |
| item:region | 1609 | 201 | 8 | 682.05 | 0.4639 | 0.8816696 |
| condition:ph | 323 | 323 | 1 | 344.98 | 0.7456 | 0.3884859 |
| condition:pressure | 232 | 232 | 1 | 466.03 | 0.5358 | 0.4645279 |
| condition:temperature | 42 | 42 | 1 | 43.75 | 0.0976 | 0.7562384 |
| item:ph | 4925 | 1231 | 4 | 682.05 | 2.8396 | 0.0235759 * |
| item:pressure | 5289 | 1322 | 4 | 682.05 | 3.0497 | 0.0165619 * |
| item:temperature | 5241 | 1310 | 4 | 682.05 | 3.0218 | 0.0173608 * |
| region:ph | 1123 | 562 | 2 | 729.83 | 1.2954 | 0.2744179 |
| region:pressure | 514 | 257 | 2 | 717.05 | 0.5926 | 0.5531316 |
| region:temperature | 1651 | 826 | 2 | 701.67 | 1.9041 | 0.1497268 |
| condition:item:region | 780 | 98 | 8 | 682.05 | 0.2250 | 0.9864123 |
| condition:item:ph | 336 | 84 | 4 | 682.05 | 0.1936 | 0.9417517 |
| condition:item:pressure | 1726 | 431 | 4 | 682.05 | 0.9951 | 0.4094523 |
| condition:item:temperature | 755 | 189 | 4 | 682.05 | 0.4353 | 0.7831263 |
| condition:region:ph | 150 | 75 | 2 | 483.89 | 0.1732 | 0.8410048 |
| condition:region:pressure | 1396 | 698 | 2 | 473.00 | 1.6100 | 0.2009734 |
| condition:region:temperature | 344 | 172 | 2 | 701.36 | 0.3964 | 0.6728689 |
| item:region:ph | 11957 | 1495 | 8 | 682.05 | 3.4472 | < 0.001 *** |
| item:region:pressure | 2255 | 282 | 8 | 682.05 | 0.6502 | 0.7354894 |
| item:region:temperature | 2145 | 268 | 8 | 682.05 | 0.6185 | 0.7627258 |
| condition:item:region:ph | 8404 | 1050 | 8 | 682.05 | 2.4228 | 0.0138468 * |
| condition:item:region:pressure | 1673 | 209 | 8 | 682.05 | 0.4822 | 0.8692024 |
| condition:item:region:temperature | 3431 | 429 | 8 | 682.05 | 0.9892 | 0.4430677 |

**Table S1**. Model 1 type II analysis of variance table with Satterthwaite's method. Blue rows show interactions that were further analyzed through post-hoc tests. Significance codes: '***' < 0.001 '**' < 0.01 '*' < 0.05

| *item* | *T.trend* | *SE* | *df* | *t* | *p* |
|---|---|---|---|---|---|
| *ownership* | −2.26 | 5.94 | 366 | −0.381 | 0.7036 |
| *agency* | 3.89 | 5.94 | 366 | 0.654 | 0.5132 |
| *location* | −7.72 | 5.94 | 366 | −1.300 | 0.1944 |
| *no body* | −13.49 | 5.94 | 366 | −2.272 | 0.0237∗ |
| *two bodies* | 2.72 | 5.94 | 366 | 0.459 | 0.6468 |

**Table S2.** Model 1 post-hoc analysis of the interaction between item and temperature (*T.trend*). Degrees-of-freedom (*df*) method: Kenward-Roger.

| *item* | *P.trend* | *SE* | *df* | *t* | *p* |
|---|---|---|---|---|---|
| *ownership* | 0.0829 | 2.25 | 710 | 0.037 | 0.9707 |
| *agency* | 2.5662 | 2.25 | 710 | 1.139 | 0.2552 |
| *location* | 5.6242 | 2.25 | 710 | 2.496 | 0.0128∗ |
| *no body* | −0.4251 | 2.25 | 710 | −0.189 | 0.8504 |
| *two bodies* | 0.3640 | 2.25 | 710 | 0.161 | 0.8718 |

**Table S3.** Model 1 post-hoc analysis of the interaction between item and pressure (*P.trend*). Degrees-of-freedom (*df*) method: Kenward-Roger. Symbols and abbreviations as in Table S2.

| *condition* | *item* | *region* | *ph.trend* | *SE* | *df* | *t* | *p* |
|---|---|---|---|---|---|---|---|
| *incongruent* | *ownership* | *stomach* | −0.899 | 6.32 | 705 | −0.142 | 0.8869 |
| *congruent* | *ownership* | *stomach* | 2.098 | 5.36 | 707 | 0.391 | 0.6959 |
| *incongruent* | *agency* | *stomach* | −10.605 | 6.32 | 705 | −1.678 | 0.0937 |
| *congruent* | *agency* | *stomach* | −1.189 | 5.36 | 707 | −0.222 | 0.8246 |
| *incongruent* | *location* | *stomach* | 21.017 | 6.32 | 705 | 3.326 | < 0.001∗∗∗ |
| *congruent* | *location* | *stomach* | 4.700 | 5.36 | 707 | 0.876 | 0.3813 |
| *incongruent* | *nobody* | *stomach* | −13.201 | 6.32 | 705 | −2.089 | 0.0370∗ |
| *congruent* | *nobody* | *stomach* | −11.393 | 5.36 | 707 | −2.124 | 0.0340∗ |
| *incongruent* | *two bodies* | *stomach* | −6.113 | 6.32 | 705 | −0.967 | 0.3337 |
| *congruent* | *two bodies* | *stomach* | −0.259 | 5.36 | 707 | −0.048 | 0.9615 |
| *incongruent* | *ownership* | *small bowel* | 16.027 | 9.52 | 705 | 1.684 | 0.0926 |
| *congruent* | *ownership* | *small bowel* | 8.706 | 9.41 | 707 | 0.925 | 0.3554 |
| *incongruent* | *agency* | *small bowel* | 24.215 | 9.52 | 705 | 2.545 | 0.0111∗ |
| *congruent* | *agency* | *small bowel* | 12.209 | 9.41 | 707 | 1.297 | 0.1950 |
| *incongruent* | *location* | *small bowel* | −8.017 | 9.52 | 705 | −0.842 | 0.3998 |
| *congruent* | *location* | *small bowel* | −0.110 | 9.41 | 707 | −0.012 | 0.9907 |
| *incongruent* | *nobody* | *small bowel* | −24.996 | 9.52 | 705 | −2.627 | 0.0088∗∗ |
| *congruent* | *nobody* | *small bowel* | 3.646 | 9.41 | 707 | 0.387 | 0.6987 |
| *incongruent* | *two bodies* | *small bowel* | 10.315 | 9.52 | 705 | 1.084 | 0.2787 |
| *congruent* | *two bodies* | *small bowel* | 6.815 | 9.41 | 707 | 0.724 | 0.4693 |
| *incongruent* | *ownership* | *large bowel* | 1.102 | 6.56 | 707 | 0.168 | 0.8666 |
| *congruent* | *ownership* | *large bowel* | 9.325 | 6.44 | 708 | 1.448 | 0.1480 |
| *incongruent* | *agency* | *large bowel* | 1.368 | 6.56 | 707 | 0.209 | 0.8348 |
| *congruent* | *agency* | *large bowel* | 7.540 | 6.44 | 708 | 1.171 | 0.2420 |
| *incongruent* | *location* | *large bowel* | −15.444 | 6.56 | 707 | −2.355 | 0.0188∗ |
| *congruent* | *location* | *large bowel* | 5.818 | 6.44 | 708 | 0.904 | 0.3665 |
| *incongruent* | *nobody* | *large bowel* | 4.836 | 6.56 | 707 | 0.737 | 0.4611 |
| *congruent* | *nobody* | *large bowel* | −0.322 | 6.44 | 708 | −0.050 | 0.9602 |
| *incongruent* | *two bodies* | *large bowel* | −1.075 | 6.56 | 707 | −0.164 | 0.8698 |
| *congruent* | *two bodies* | *large bowel* | −10.558 | 6.44 | 708 | −1.640 | 0.1015 |

**Table S4**. Model 1 post-hoc spotlight analysis of the interaction between condition, item, region, and pH (*ph.trend*). Symbols and abbreviations as in Table S2. Degrees-of-freedom (*df*) method: Kenward-Roger.

| | *Sum Sq* | *Mean Sq* | *NumDF* | *DenDF* | *F* | *p* |
|---|---|---|---|---|---|---|
| condition | 33017 | 33017 | 1 | 228.72 | 67.7224 | $< 0.001$ *** |
| item | 33277 | 8319 | 4 | 213.57 | 17.0637 | $< 0.001$ *** |
| EGG peak | 1303 | 1303 | 1 | 172.15 | 2.6719 | 0.10396 |
| condition:item | 30907 | 7727 | 4 | 213.57 | 15.8484 | $< 0.001$ *** |
| condition:EGG peak | 2246 | 2246 | 1 | 182.29 | 4.6064 | 0.03317 * |
| item:EGG peak | 1270 | 318 | 4 | 213.57 | 0.6515 | 0.62645 |
| condition:item:EGG peak | 880 | 220 | 4 | 213.57 | 0.4512 | 0.77142 |

**Table S5**. Model 2 type II analysis of variance table with Satterthwaite's method. The blue row shows the interaction that was further analyzed through post-hoc tests. Significance codes as in table S1.

| *condition* | *EGG.trend* | *SE* | *df* | *t* | *p* |
|---|---|---|---|---|---|
| *incongruent* | −293 | 463 | 190 | −0.634 | 0.5268 |
| *congruent* | 972 | 384 | 161 | 2.534 | 0.0122* |

**Table S6**. Model 2 post-hoc analysis of the interaction between condition and EGG peak frequency (*EGG.trend*). Symbols and abbreviations as in Table S2. Degrees-of-freedom (*df*) method: Kenward-Roger.

| | *interoception scores* | | |
|---|---|---|---|
| | sensibility | accuracy | awareness |
| *N* | 27 | 31 | 31 |
| *mean* | 2.534 | 0.669 | 50.124 |
| *SD* | 0.945 | 0.202 | 23.869 |
| *min* | 0.800 | 0.140 | 0 |
| *max* | 4.390 | 0.963 | 90.909 |

**Table S7**. Descriptive statistics of interoceptive sensibility (scale: 0-5), accuracy (scale: 0-1), and awareness (scale: 0-100).

**Movie S1.**

Movie describing each phase of the embreathment illusion. Available at https://youtu.be/4zBx27OoIRE
